# Intact early visual representations, not phosphene-adapted features, account for human perceptual behavior with retinal prostheses

**DOI:** 10.1101/2025.06.23.660990

**Authors:** Jonathan Skaza, Shravan Murlidaran, Apurv Varshney, Ziqi Wen, Miguel P. Eckstein, Michael Beyeler

**Affiliations:** Graduate Program in Dynamical Neuroscience, University of California, Santa Barbara; Department of Psychological and Brain Sciences, University of California, Santa Barbara; Department of Computer Science, University of California, Santa Barbara

## Abstract

Efforts to restore vision via neural implants have outpaced the ability to predict what users will perceive, leaving patients and clinicians without reliable tools for surgical planning or device selection. To bridge this critical gap, we introduce a computational virtual patient (CVP) pipeline that integrates anatomically grounded phosphene simulation with task-optimized deep neural networks (DNNs) to forecast patient perceptual capabilities across diverse prosthetic designs and tasks. We evaluate performance across six visual tasks, six electrode configurations, and two artificial vision models, establishing our CVP approach as a scalable pre-implantation method. Several chosen tasks align with the Functional Low-Vision Observer Rated Assessment (FLORA), revealing correspondence between model-predicted difficulty and real-world patient outcomes. Further, the CVP paradigm exhibited strong correspondence with psychophysical data collected from normally sighted subjects viewing phosphene simulations, capturing both overall task difficulty and performance variation across implant configurations. Comparing frozen-feature linear probing with full end-to-end fine-tuning reveals that preserving natural-image representations—rather than adapting them to phosphene-specific statistics—better reproduces human perceptual behavior, consistent with the constrained plasticity of adult visual cortex. The findings position CVP as a scientific tool for probing perception under prosthetic vision, an engine to inform device development, and a clinically relevant framework for pre-surgical forecasting.

---

Restoring sight to individuals with severe vision loss remains one of the foremost challenges in neuroscience and bioengineering. Among the leading causes of irreversible blindness is retinitis pigmentosa (RP), a group of inherited retinal degenerations for which gene and stem-cell therapies show promise only in early stages [1, 2, 3]. For those with advanced retinal or optic nerve damage, electronic visual prostheses (*bionic eyes*) offer a technological alternative by electrically stimulating surviving neurons to evoke visual percepts known as *phosphenes* [4, 5]. Modern devices combine an external camera, a vision processing unit that converts images into electrical stimulation patterns, and an electrode array implanted along the visual pathway: either in the retina [6, 7, 8], optic nerve [9], thalamus [10], or visual cortex [11, 12, 13].

Long-term studies of retinal implants, such as Argus II [6] and PRIMA [7], and cortical implants, such as Orion [14], ICVP [12], and CORTIVIS [11], confirm device safety and stability over many years [15, 16, 17, 8]. Yet the percepts they evoke remain coarse and distorted [18, 19, 20, 21], supporting only basic visual functions such as light detection, motion perception, and high-contrast object localization [22, 23, 8, 24]. PRIMA is a notable exception, enabling limited reading performance in geographic atrophy [25]. Across users, outcomes vary widely due to differences in residual retinal health [26, 27, 28], cortical plasticity [29, 30, 31], and visual training [32, 33, 34, 35].

Clinical translation remains slow and expensive. Pivotal trials for Argus II and PRIMA have required years of engineering refinement, surgical implantation, and longitudinal testing in small cohorts. Standardized instruments such as the Functional Low-Vision Observer Rated Assessment (FLORA) [36] represent important steps toward quantifying real-world performance by measuring patients’ ability to navigate, recognize people, and locate objects, but these assessments still depend on trained observers, site-specific setups, and subjective ratings. Each new device iteration or algorithmic update necessitates a fresh round of clinical evaluation, constraining innovation and slowing progress. This imbalance between rapid technological development and the pace of human testing underscores the need for predictive, scalable models that can estimate functional outcomes before surgery.

To complement such slow and costly testing, researchers have long relied on *simulated prosthetic vision (SPV)* experiments with sighted observers to study artificial vision under controlled conditions. Early work displayed phosphene-like patterns on monitors to probe letter, face, and object recognition [37, 38, 39], while more recent studies have used immersive or augmented-reality paradigms that incorporate head and eye movements in naturalistic environments [40, 41, 42, 43]. These paradigms have revealed scanning and gaze strategies resembling those of prosthesis users during reading [44], quantified trade-offs between spatial resolution and field of view [45, 46], and shown how phosphene density and shape influence navigation and obstacle avoidance [47, 48, 49, 41]. Further work has examined perceptual learning [50, 34] and tested scene-simplification strategies to enhance recognition and mobility [51, 52, 53, 54, 55]. Collectively, these studies establish SPV as a robust experimental framework for linking stimulation parameters to visual behavior and for probing the limits of artificial vision without the need for implantation.

A natural next step is to move beyond isolated experiments toward predictive models of prosthetic vision. A *computational virtual patient* [56] (i.e., a computational model integrating anatomy, stimulation, and behavior) aims not only to reproduce phosphene appearance but to predict behavioral performance across devices and tasks. Comparable *in silico* clinical trial frameworks are already transforming other areas of medicine, where individualized simulations are used to test and optimize drugs, cardiac implants, medical image quality [57, 58, 59, 60], and neuromodulation therapies before human trials [61, 62]. In prosthetic vision, existing phosphene models can accurately map stimulation parameters to perceptual outcomes when constrained by neuroanatomical, neurophysiological, or psychophysical data [18, 63, 64, 65, 66]. Yet these models remain primarily descriptive: they simulate percepts under known conditions rather than predicting how those percepts drive functional performance.

A computational virtual patient must go further, embedding mechanistic models of perception within a behavioral framework that predicts human performance across tasks. Two recent studies by An *et al*. explored this idea using low-resolution phosphene simulations of faces: one applied supervised learning to predict match-to-sample outcomes [67], while the other used reinforcement learning to emulate learning dynamics across trials [68]. Although these studies showed that artificial agents can approximate human performance, they were restricted to a single task under pixelated vision and lacked biological realism. No existing framework yet predicts behavioral performance across implant types, stimulation models, and functional tasks within a unified, physiologically grounded model.

To address this gap, we present a *computational virtual patient (CVP)* framework that operationalizes the concept of a virtual patient for prosthetic vision (Figure 1). The framework integrates anatomically grounded phosphene simulations with deep neural networks (DNNs) trained to perform ecologically relevant tasks such as object recognition and spatial localization. To investigate whether task performance relies on reinterpretation of existing natural-image representations or requires adaptation to phosphene-specific statistics, we compare frozen-feature linear probing with full end-to-end fine-tuning and evaluate which better reproduces human perceptual behavior. Linear probing holds low-level features fixed while adapting only downstream readout—a computational analogue of the constrained plasticity observed in adult prosthesis users, whose pre-existing visual circuitry remains largely intact and whose learning reflects reinterpretation of degraded input rather than reorganization of early visual representations [69, 70]. We validate the CVP’s predictions against psychophysical data from sighted participants and FLORA clinical performance metrics from implanted users. The framework accurately reproduces human performance and device-dependent trends, establishing computational virtual patients as scalable surrogates for behavioral testing and offering a principled path to forecast functional vision, accelerate device optimization, and set realistic expectations for future recipients of visual prostheses.

**Figure 1:**
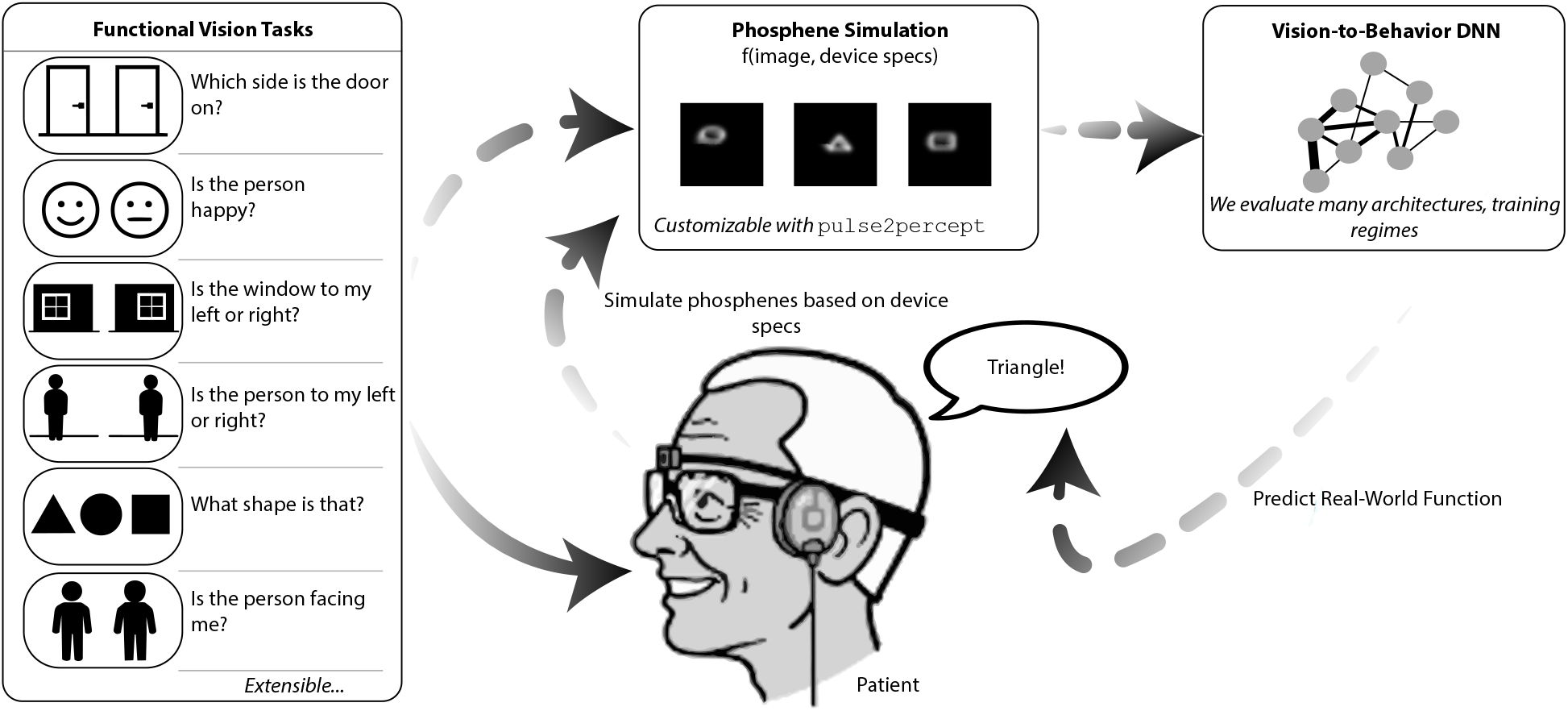
Overview of the computational virtual patient (CVP) pipeline. The framework comprises three stages: (i) generation of real-world functional vision task stimuli; (ii) prosthetic vision simulation via device-specific phosphene rendering; and (iii) task evaluation using vision-to-behavior deep neural networks (DNNs). Together, these components enable systematic comparison across electrode configurations, artificial vision models, allowing scalable, pre-implantation prediction of real-world functional performance.

## Results

### Overview of the computational virtual patient framework

The CVP framework reproduces core dependencies observed in psychophysical and clinical studies of prosthetic vision by integrating three stages: (i) generation of ecologically relevant stimuli, (ii) simulation of prosthetic vision using biophysically informed phosphene models, and (iii) behavioral evaluation by computational observers (Figure 2). This task-agnostic pipeline can, in principle, be applied to any functional vision task by transforming naturalistic inputs into device-specific phosphene representations and evaluating downstream performance.

**Figure 2:**
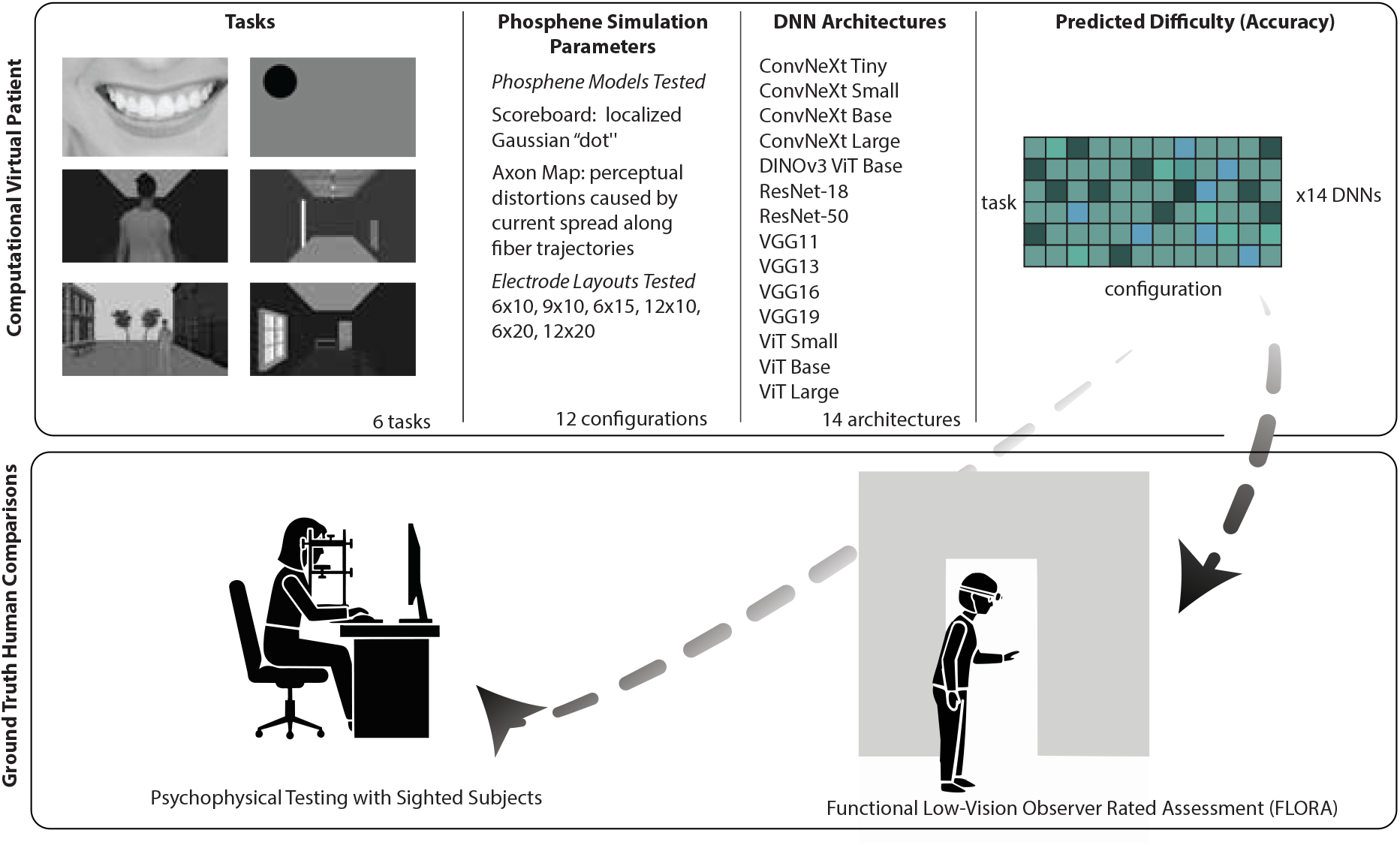
Validation of the CVP pipeline through psychophysical and clinical comparisons. We simulate stimuli for six classification tasks and transform the raw inputs into phosphene representations using 12 prosthetic device parameterizations. Task performance is then compared between various DNNs and humans, using psychophysical evaluations on identical stimuli and real-world subjective outcomes reported by prosthesis users.

To demonstrate this approach, we instantiated six classification tasks reflecting everyday perceptual challenges encountered by prosthesis users [36, 24]: emotion identification, shape classification, direction a person is facing, and spatial localization of doors, people, and windows (Table 1). These tasks span a range of perceptual difficulty and include activities assessed in the FLORA, enabling comparison between simulated task difficulty and subjective ratings reported by implanted users. Eighteen sighted participants additionally performed three of these tasks (emotion identification, shape classification, doorway localization) on phosphene-rendered stimuli, providing direct psychophysical benchmarks (see Methods for experimental details).

**Table 1:**
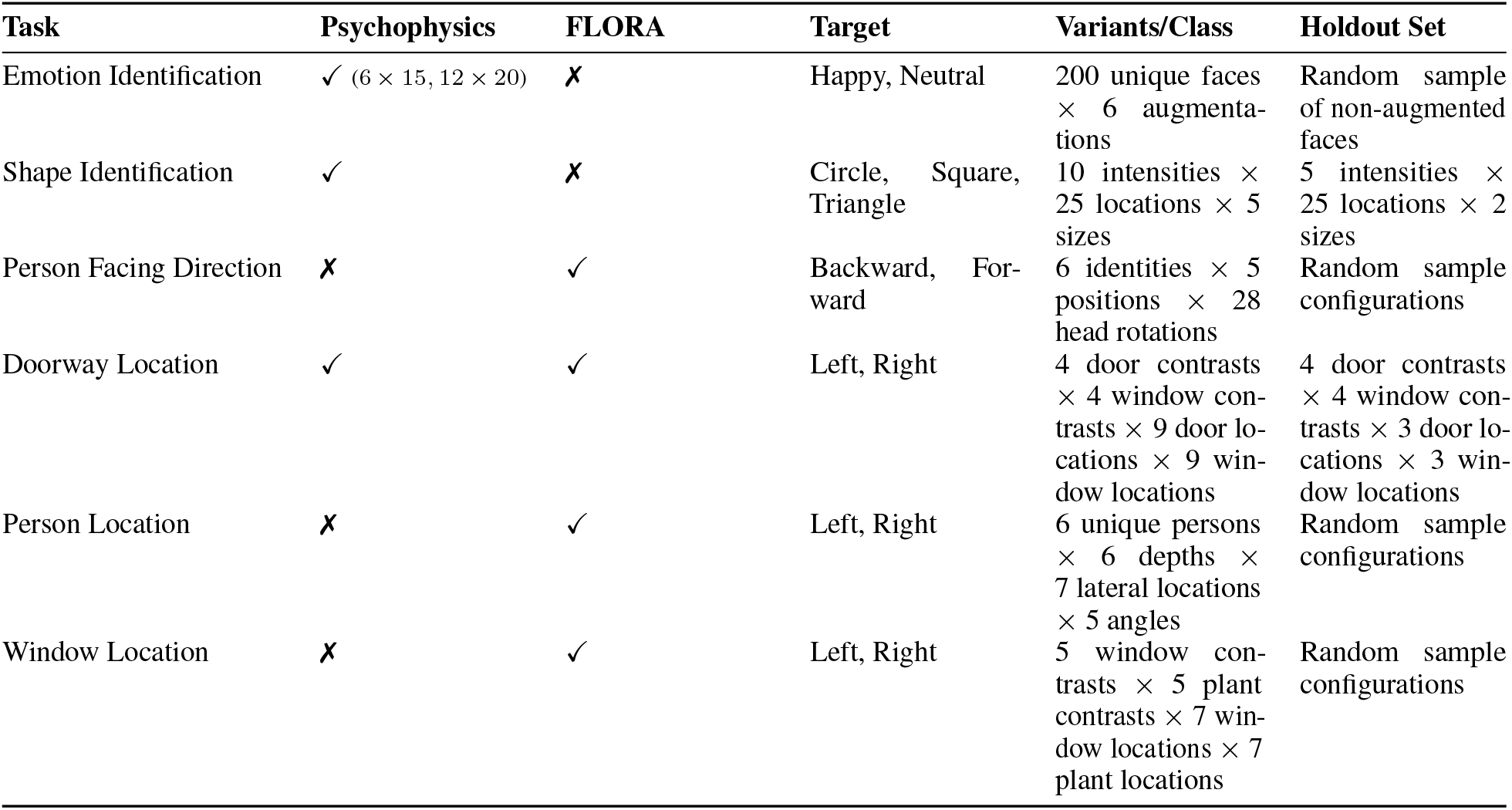
Overview of tasks, experimental inclusion, and stimulus parameterizations. The **Psychophysics** and **FLORA** columns indicate whether each task was included in human behavioral testing and whether an analogous task was implemented in the FLORA, respectively. The **Target** column lists the classification labels for each task. The **Variants/Class** column specifies the full combinatorial parameter space used to generate task stimuli. The **Holdout Set** column describes the subset of parameter combinations reserved for evaluation, enabling assessment of human and/or model performance.

| Task | Psychophysics | FLORA | Target | Variants/Class | Holdout Set |
| --- | --- | --- | --- | --- | --- |
| Emotion Identification | ✓ (6 × 15, 12 × 20) | ✗ | Happy, Neutral | 200 unique faces × 6 augmentations | Random sample of non-augmented faces |
| Shape Identification | ✓ | ✗ | Circle, Square, Triangle | 10 intensities × 25 locations × 5 sizes | 5 intensities × 25 locations × 2 sizes |
| Person Facing Direction | ✗ | ✓ | Backward, Forward | 6 identities × 5 positions × 28 head rotations | Random sample configurations |
| Doorway Location | ✓ | ✓ | Left, Right | 4 door contrasts × 4 window contrasts × 9 door locations × 9 window locations | 4 door contrasts × 4 window contrasts × 3 door locations × 3 window locations |
| Person Location | ✗ | ✓ | Left, Right | 6 unique persons × 6 depths × 7 lateral locations × 5 angles | Random sample configurations |
| Window Location | ✗ | ✓ | Left, Right | 5 window contrasts × 5 plant contrasts × 7 window locations × 7 plant locations | Random sample configurations |

Stimuli were transformed using two established phosphene models implemented in pulse2percept [71]. The scoreboard model, which approximates subretinal and cortical stimulation [72, 64], produced localized Gaussian “dots”, whereas the axon map model, designed for epiretinal implants, incorporated current spread along retinal ganglion cell axons to produce elongated, comet-shaped percepts [18, 63] (examples in Figure S1). These morphologies were combined with six electrode layouts spanning and extending the Argus II geometry (6 × 10, 9 × 10, 6 × 15, 12 × 10, 6 × 20, 12 × 20), systematically varying spatial density and coverage.

To examine the impact of DNN backbone on task performance and human alignment under phosphene stimulation, we conducted a large-scale evaluation spanning six tasks, two phosphene models, six electrode configurations, and 14 distinct architectures—including convolutional neural networks (CNNs) and vision transformers (ViTs)—encompassing VGG [73], ResNet [74], ConvNeXt [75, 76], ViT [77], and DINOv3 [78]. For each task–configuration pair, we trained 10 independently initialized models (random seeds) per architecture to obtain stable performance estimates. In this section, we focus on models trained using a linear probe approach, in which pretrained backbones were frozen and only task-specific linear heads were optimized. This isolates how well natural-image representations support functional vision under phosphene stimulation without additional feature adaptation. Full end-to-end fine-tuning of the backbone is explored in a subsequent subsection. Parallel comparison of human and model behavior using overall task performance, trends across implant configurations, and trial-level error patterns establishes the CVP as a scalable platform for *in silico* psychophysics linking controlled laboratory benchmarks with clinically reported functional outcomes.

### Alignment in difficulty across CVPs and human observers

Computational virtual patients reproduced human perceptual behavior with high fidelity across tested conditions. While models achieved slightly higher absolute accuracy than human observers, both preserved identical task difficulty rankings and exhibited comparable variance across electrode configurations and phosphene simulation models. Quantifying this correspondence, we computed correlations between normalized above-chance performance (to account for different chance levels across tasks, see Methods) in humans and CVPs across the 28 psychophysical conditions tested, revealing strong and statistically significant alignment across all backbone architectures, ranging from *r* = 0.874 (*p <* .001) for ConvNeXt Base to *r* = 0.941 (*p <* .001) for VGG19.

This alignment was present across multiple dimensions of perceptual performance. First, CVPs (aggregated across architectures and configurations) and humans maintained the same relative ordering of task difficulty across all six tested scenarios: emotion identification, shape discrimination, and spatial localization (Figure 3). Critically, for FLORA-inspired tasks lacking direct psychophysical task-performance benchmarks, CVP performance aligned with subjective difficulty ratings reported by prosthesis users (Figure 3), indicating that the framework extends beyond controlled laboratory conditions to capture real-world clinical outcomes.

**Figure 3:**
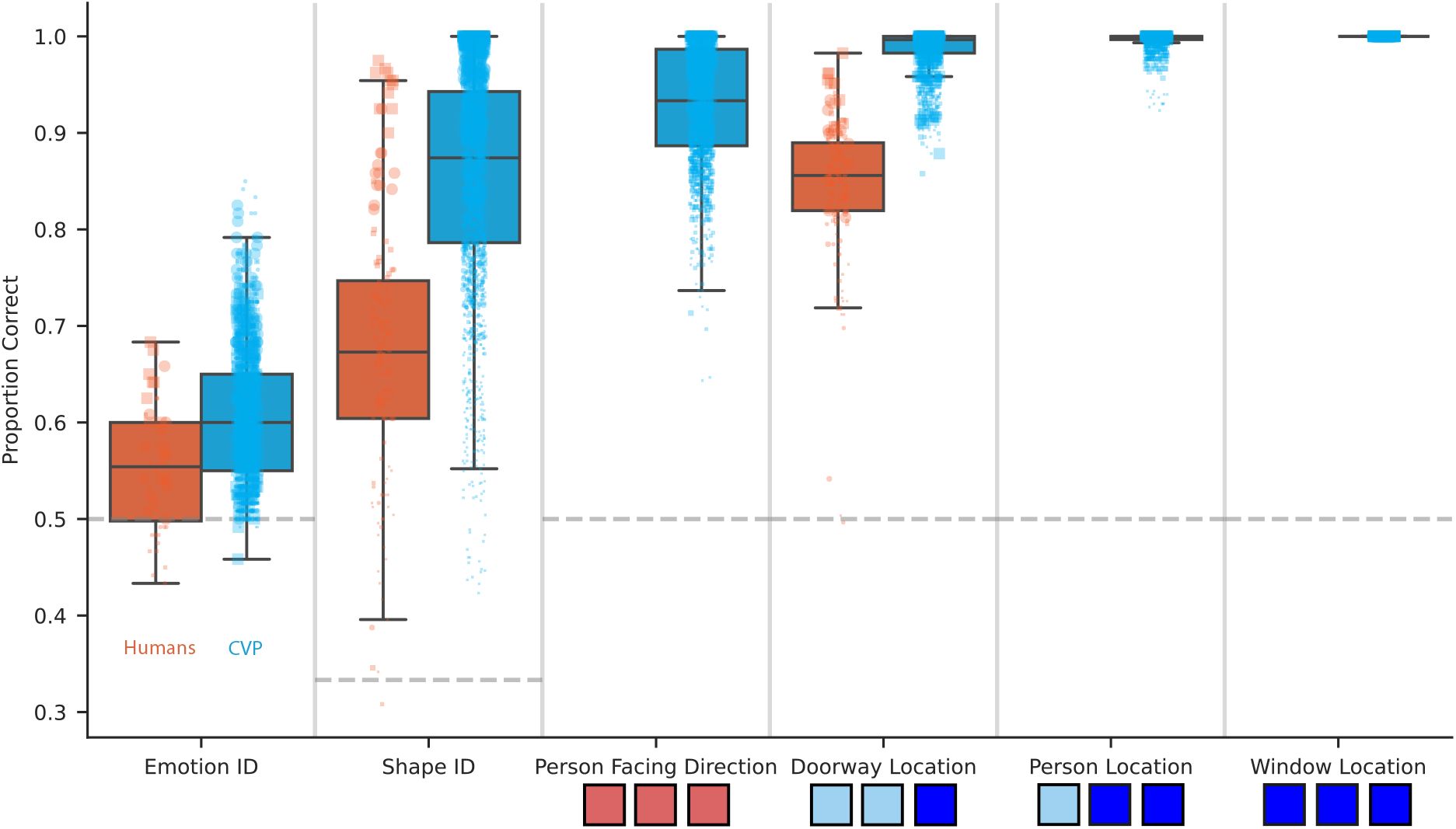
Performance of human and artificial observers across tasks. Performance across six simulated bionic vision tasks, comparing human observers (orange) and CVPs using various DNN backbones (blue). Boxplots summarize the distribution of proportion correct (*PC*) scores: the line inside each box represents the median (50^*th*^ percentile); the bottom and top edges of the box correspond to the first (Q1, 25^*th*^ percentile) and third quartiles (Q3, 75^*th*^ percentile), respectively, with the box height representing the interquartile range (*IQR*). Whiskers extend to the most extreme data points within 1.5 × *IQR* from the box edges. Overlaid points represent individual human observers or model instances, with marker area proportional to the number of electrodes (60 → 240) and shape indicating the phosphene simulation (square = scoreboard; circle = axon map). Dashed horizontal lines denote chance-level performance for the task. Tasks are ordered by mean accuracy to highlight relative difficulty. Below select task labels, FLORA-based clinical difficulty ratings from three prosthesis users are shown using color codes (red = difficult, light blue = moderate, dark blue = easy), based on ease of task scores at the final available study endpoint [24]. The human-perceived difficulty ordering is preserved in CVP performance trends, suggesting shared sensitivity to task demands.

Second, consistent with spatial sampling theory, both humans and CVPs exhibited systematic performance scaling with electrode count, and this held across both phosphene models (Figure 4). Task performance increased monotonically from the 6 × 10 configuration (60 electrodes) through the 6 × 15 baseline (90 electrodes) to the 12× 20 high-density array (240 electrodes) across most task-phosphene combinations. However, the magnitude of these gains varied substantially by task. Shape identification exhibited the greatest sensitivity to electrode density in both humans and CVPs, with performance improvements consistently outpacing those observed for doorway location and emotion identification. For the shape task under the scoreboard phosphene model, human performance improved by Δ*PC* = 0.426 between the lowest and highest density configurations, while CVPs showed Δ*PC* = 0.360.

**Figure 4:**
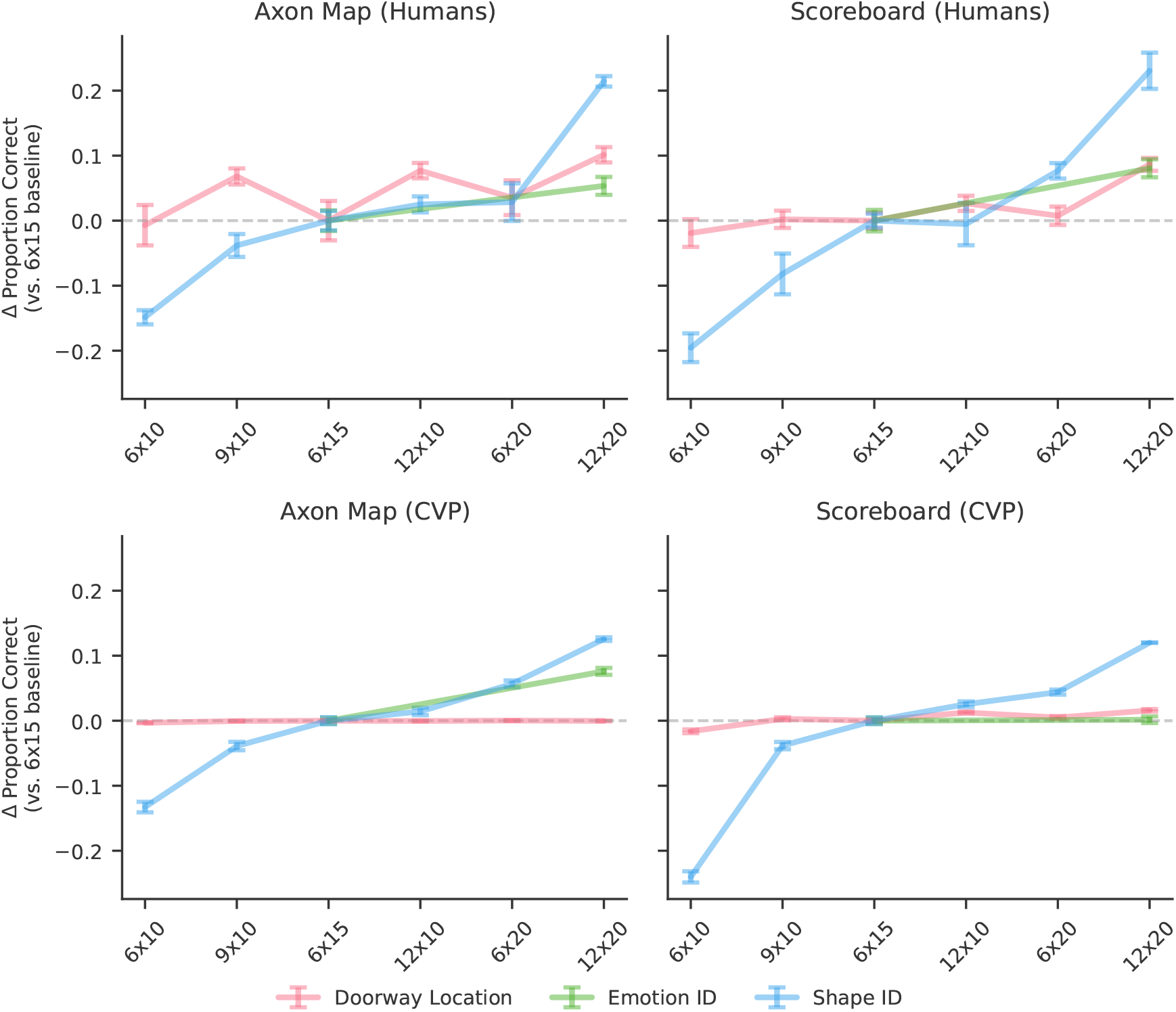
Change in proportion correct (Δ*PC*) as a function of implant resolution, phosphene model, and task. Performance is expressed relative to the 6 × 15 implant baseline (dashed line) within each task and phosphene model. Columns compare phosphene generation models (axon map vs. scoreboard), and rows compare human participants (top) with computational virtual patients (CVPs; bottom). The *x*-axis denotes electrode electrode array resolution. Colored lines show task-specific mean Δ*PC* across implant configurations for doorway location, emotion identification, and shape identification; error bars indicate SEM across subjects (humans) or independent model runs (CVPs).

Third, device-specific perceptual effects exhibited similar attenuation across humans and CVPs, indicating that CVPs can capture not only overall performance levels but also the qualitative impact of phosphene-induced distortions. In the shape identification task—where performance varied most strongly across electrode configurations—we compared performance across the 6 ×10 and 12 ×20 arrays under the two phosphene models: the scoreboard model and the axon map. Under the axon map rendering, both humans and CVPs showed a compressed performance range between these array resolutions relative to the scoreboard condition. In humans, this range decreased from 0.426 (scoreboard) to 0.363 (axon map), while in CVPs it decreased from 0.360 to 0.258. This similar attenuation suggests that computational virtual patients are sensitive to the same axonal distortions that constrain effective spatial resolution in human prosthetic vision.

### Error pattern analysis between humans and computational virtual patients reveals architectural differences

Beyond aggregate performance metrics, we examined whether humans and CVPs exhibited similar patterns of trial-level errors—a more fine-grained test of perceptual alignment. For each DNN architecture, we computed the Jaccard similarity index between the sets of trials that elicited errors in human psychophysics versus models across all shared task–configuration pairs. The Jaccard index quantifies the overlap between two error sets as *J* (*A, B*) = |*A* ∩ *B*| */* |*A* ∪ *B* |, where values near 0 indicate no overlap and values near 1 indicate near-complete agreement. Similarity indices were calculated for each (human subject, model run) pair and averaged using weights proportional to the number of common trials to obtain a similarity score for each model architecture (Figure 5). As an upper-bound reference, we first computed human–human error similarity across all subject pairs, yielding a Jaccard index of 0.284 ± 0.016 (bootstrap SE), providing a noise ceiling for the maximum alignment one could expect given inter-observer variability.

**Figure 5:**
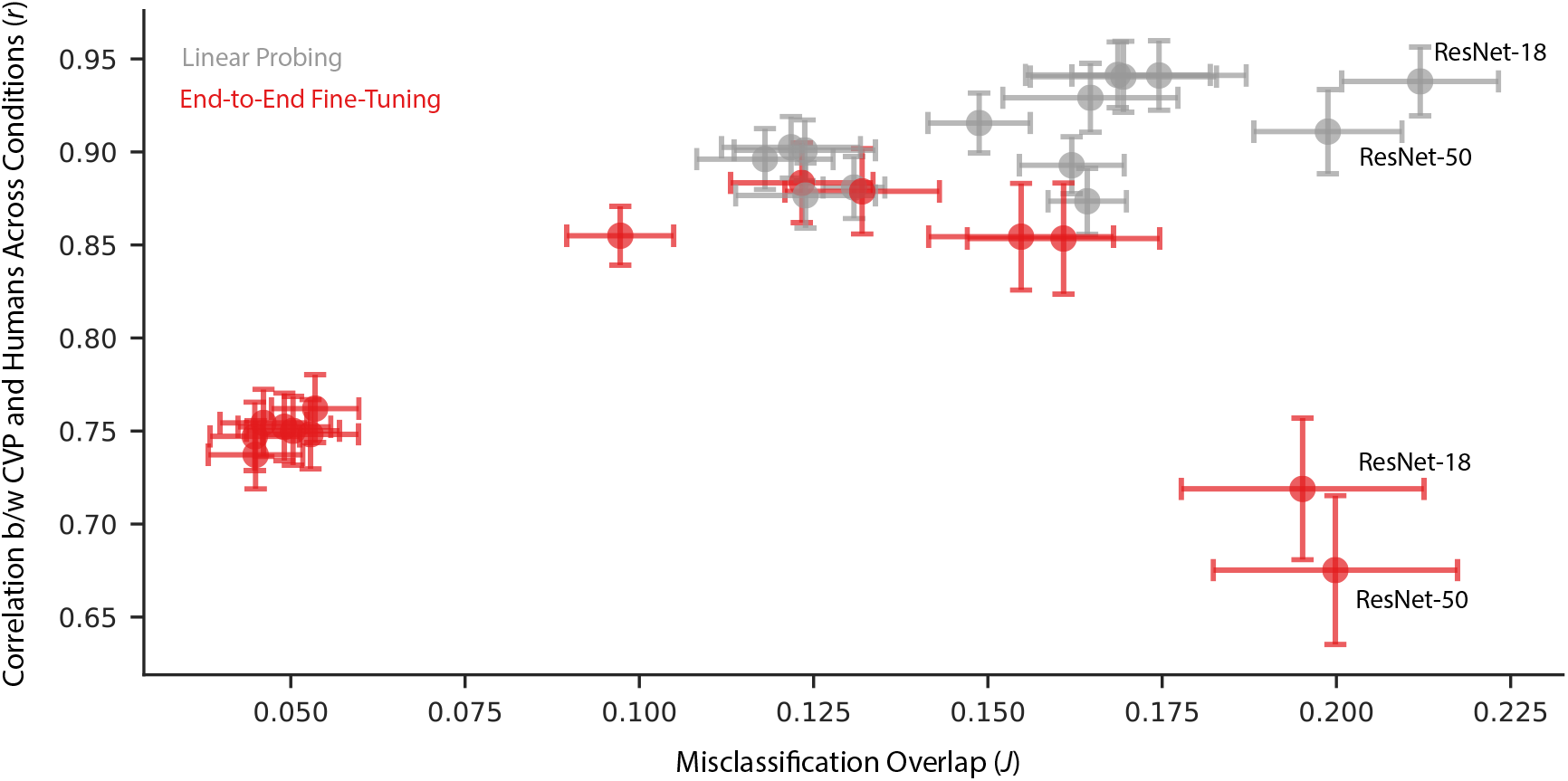
Relationship between two measures of model–human similarity across DNN backbones. Each point represents one architecture trained with either linear probing (gray) or full fine-tuning (red). The *x*-axis shows the Jaccard index of misclassification overlap—the proportion of shared errors between model and human observers on matched trials. The *y*-axis shows the Pearson correlation between model and human condition-varying performance. Error bars correspond to bootstrapped SE (subject-level resampling) for each metric. ResNets are annotated as an interesting case that has high error overlap regardless of training scheme.

As above, we focused our primary analysis on computational virtual patients trained via linear probing—that is, a linear classifier trained on frozen pretrained features. Across architectures, this method produced reliable and significantly above chance alignment with human error patterns (permutation test, *p <* 0.001 for all architectures, complete results across all architectures and training regimes are reported in Table S1.), demonstrating that the internal representations of pretrained DNNs systematically encode stimulus-level difficulty in ways that partially mirror human perception.

Model–human Jaccard indices under linear probing ranged from *J* = 0.118 ± 0.010 (DINOv3 ViT Base) to *J* = 0.212 ± 0.011 (ResNet-18). The architecture family was a strong determinant of alignment. Convolutional networks— ResNets and VGG-family models—showed the highest human–model error concordance: ResNet18 achieved the best alignment (*J* = 0.212 ± 0.011), followed closely by ResNet50 (*J* = 0.199 ± 0.011) and VGG-family networks (*J* ≈ 0.165–0.174). ConvNeXt architectures, which blend convolutional inductive biases with modernized design choices, fell at an intermediate level (*J* ≈ 0.130–0.162), with performance varying across model scale.

In contrast, Vision Transformer (ViT) architectures consistently showed the weakest alignment with human error patterns, with all ViT variants clustering near *J* ≈ 0.12—regardless of model size (ViT Small: *J* = 0.124 ± 0.010; ViT Base: *J* = 0.122 ± 0.010; ViT Large: *J* = 0.124± 0.010). DINOv3 ViT Base, despite being trained with a self-supervised objective that promotes richer semantic representations, similarly yielded low alignment in this analysis (*J* = 0.118 ± 0.010). The convergence of ViT models near a common, lower bound—independent of scale or pretraining strategy—suggests that the transformer architecture’s lack of explicit local spatial inductive biases may fundamentally limit its alignment with human perceptual failures, at least when representations are read out linearly.

### End-to-end fine-tuning degrades human–model alignment

Full end-to-end fine-tuning (i.e., training all backbone weights on phosphene-rendered stimuli) resulted in worse perceptual alignment compared to linear probing. Mean Pearson correlation across architectures dropped from *r* = 0.910 (linear probe) to *r* = 0.784 (end-to-end); bootstrap tests confirmed linear probing was significantly better aligned for all architectures (all *p <* 0.001; Table S3). Trial-level Jaccard indices similarly declined. ResNets were a notable exception, maintaining high error overlap under both regimes (Figure 5). Gaussian noise injection applied to fully fine-tuned logits recovered aggregate accuracy but did not restore condition-level alignment to linear probe levels (additive: *r* = 0.870; proportional: *r* = 0.878; both *p <* 0.001 vs. linear probe; Tables S2, S3), indicating that fine-tuning introduces representational distortions not correctable at the output level.

Taken together, these results indicate that computational virtual patients based on convolutional feature extractors capture stimulus-specific failure patterns that more closely mirror those of human observers. That alignment was recoverable via simple linear probing—without any fine-tuning of the backbone—further suggests that the relevant perceptual geometry is already encoded in the frozen representations of these networks. The gap that remains relative to the human–human noise ceiling (*J* = 0.284) indicates that even the best-aligned architectures reproduce human perceptual behavior through partially distinct internal representations (consistent with [67]), motivating further investigation into how architectural inductive biases shape alignment with biological vision.

## Discussion

### A scalable framework for predicting functional vision

The CVP framework introduced here predicts functional vision outcomes across devices and tasks by aligning simulated phosphene vision with human psychophysics and clinical benchmarks. Traditional evaluation of prosthetic vision depends on small, long-term clinical trials such as Argus II and PRIMAvera that can take years to complete and cost millions of dollars [25, 6]. These trials have shown that while prostheses can restore basic light perception and object localization, individual outcomes vary widely [24]. The CVP addresses this bottleneck by offering a quantitative, pre-implantation approach to forecasting behavioral performance. Trained on realistic phosphene renderings, DNNs reproduced human psychophysical data and mirrored trends observed in implanted users, including improved accuracy with denser electrode arrays and hindered performance under axonal distortions [18, 79, 19]. Consistency across psychophysical, simulated, and clinical scales suggests that CVP can serve as a reliable *in silico* proxy for functional vision testing.

Structurally, CVP mirrors a training paradigm: by evaluating performance across diverse tasks, electrode arrays, and phosphene models, it establishes a foundation for generalization across users and contexts. This builds on recent work showing that DNNs can approximate human behavior in low-resolution prosthetic vision tasks [67], and complements broader evidence that DNNs can serve as computational models of human vision [80, 81, 82]. In this sense, CVP functions not merely as an engineering tool but as a framework for probing how prosthetic vision shapes perception and behavior. While not a replacement for clinical trials, it provides a scalable, standardized means of generating early-stage insights that would be costly or impractical to obtain from patient testing alone.

### Linear probing as a model of constrained perceptual adaptation

The alignment advantage of linear probing is consistent with adult visual plasticity research suggesting that prosthesis users adapt primarily by reinterpreting degraded input through largely intact pre-existing visual representations, rather than overhauling feature processing [69, 70]. Full fine-tuning, which reshapes the entire feature hierarchy around phosphene-specific statistics, may better approximate the representational flexibility available during developmental critical periods—suggesting that a CVP built from a fully fine-tuned or from-scratch network might more accurately model perceptual experience in individuals implanted in early childhood.

The linear readout result is perhaps unsurprising for our normally sighted psychophysical observers. Having viewed phosphene simulations for only a few hours across the experiment, there is no reason to expect any reorganization of their early visual processing areas; they necessarily interpret the degraded stimuli through pre-existing natural-image representations, making linear probing the expected computational analogue of their perceptual strategy. The more consequential question is whether this model also generalizes to blind patients with prosthetic implants. Evidence suggests it may, at least for those implanted beyond the critical period. Such patients show only limited perceptual learning within the first days to weeks following implantation [29], consistent with visual representations remaining largely stable and adaptation occurring primarily through downstream readout. Patients implanted during the critical period, however, may undergo more extensive reorganization of early visual cortex in response to prosthetic input, potentially rendering a CVP with greater layer-wise retraining a more faithful model of their perceptual experience.

### Human–model alignment depends on architectural inductive biases

Although CVPs matched the overall difficulty ordering observed in human psychophysics, finer-grained error analyses revealed that equivalent aggregate accuracy can arise from qualitatively different perceptual strategies, with systematic differences emerging across architectural families.

Convolutional architectures—including ResNet and VGG-family networks—showed the strongest trial-level alignment with human error patterns, as measured by the Jaccard similarity index between model and human misclassifications. Across these models, errors tended to occur on the same trials that elicited human misclassifications, indicating higher overlap in stimulus-level failure modes.

Vision Transformers consistently clustered near the lower bound of human–model alignment regardless of model scale or pretraining objective, with all ViT variants yielding Jaccard indices near *J* ≈ 0.12. This convergence was observed across architecturally distinct implementations evaluated under the same linear probing protocol. ConvNeXt architectures occupied an intermediate position between convolutional networks and transformers on this measure.

All models were evaluated using frozen pretrained representations with linear readout, and these architectural differences emerged without task-specific end-to-end fine-tuning. Notably, alignment for all model classes remained below the human–human noise ceiling (*J* = 0.284), indicating that even the most human-aligned models reproduced behavioral errors with only partial overlap at the trial level [67].

### From simulation to translation

While powerful, the current implementation is limited to static, two-dimensional stimuli and visual-only input. Real-world prosthetic vision is dynamic, embodied, and multisensory: patients integrate auditory, tactile, and proprioceptive cues while using head and eye movements to accumulate information over time [41, 33, 83]. Simulations also assume stable electrode performance and consistent percepts, whereas clinical data reveal substantial variability and adaptation [29, 26]. Future iterations of CVP should incorporate temporal sequences, multimodal inputs, and subject-specific anatomy to capture individual variability and learning. Such extensions would transform CVP from a population-level predictor into a personalized clinical tool, aligning with ongoing efforts toward patient-specific digital twins in neuroengineering [84, 65, 61].

Even in its current form, the CVP represents a shift in how prosthetic vision can be evaluated. By providing standardized, scalable, and anatomically informed behavioral predictions, it can help clinicians set expectations, guide device design, and inform regulatory review. More broadly, it exemplifies how *in silico* frameworks can bridge neuroscience and computation, accelerating translation from algorithmic innovation to tangible improvements in quality of life for people who are blind.

## Methods

We developed a CVP framework to evaluate human–model alignment in prosthetic vision. The approach integrates ecologically relevant visual tasks, biophysically informed phosphene simulations, and both human and DNN observers tested under identical image conditions. This section details task design, phosphene simulation, psychophysical testing, model training, and analysis procedures.

### Task design and stimulus generation

We investigated six simulated tasks reflecting functional challenges reported by prosthesis users [24]. The tasks were designed to span a range of visual functions and difficulty levels while remaining computationally tractable for large-scale modeling. Some tasks were adapted from the FLORA but were not administered to sighted participants, allowing comparison to published clinical outcomes.

Each task comprised many stimulus variants, systematically varying parameters such as position, intensity, contrast, and size. This yielded large and diverse datasets for model training and psychophysical evaluation. All stimuli were converted to grayscale to match the limited luminance range of prosthetic vision and were tested under 12 distinct simulation configurations. These configurations included two phosphene models (scoreboard and axon map) and six electrode layouts: 6 × 10, 9 × 10, 6 × 15, 12 × 10, 6 × 20, 12 × 20.

Stimuli for the shape task were generated using custom Python scripts, while the emotion identification task used the AffectNet dataset [85]. The remaining tasks (person facing direction, doorway location, person location, and window location) were rendered in Unity to produce naturalistic 3D scenes. All images were then processed with the `pulse2percept` package [71] to simulate prosthetic percepts.

### Phosphene simulation and device configurations

Each image was transformed into a simulated phosphene representation using one of two well-established models implemented in `pulse2percept` [71]. The scoreboard model assumes each electrode produces a localized Gaussian “dot”, providing a simplified approximation of prosthetic vision. The axon map model incorporates the geometry of retinal ganglion cell axon pathways, capturing perceptual distortions caused by current spread along fiber trajectories [18].

Stimuli were rendered for six electrode array geometries (6 × 10, 9 × 10, 6 × 15, 12 × 10, 6 × 20, 12 × 20), spanning configurations similar to commercial and experimental implants such as Argus II and PRIMA. This design allowed systematic exploration of how spatial sampling and axonal distortion influence perceptual outcomes.

### Human psychophysics

Behavioral data were collected from 18 undergraduate participants (ages 18–22) with normal or corrected-to-normal vision. All procedures were approved by the University of California Institutional Review Board.

Participants completed subsets of three tasks (emotion identification, shape classification, and doorway localization) under forced fixation. Each trial presented a phosphene-rendered image for 500 ms, followed by a response window. Subjects were trained with ten practice trials before each block.

Each condition included a subset (*n* = 12) drawn from the full participant pool (*N* = 18). Shape and doorway tasks were tested under all 12 phosphene configurations (240 and 288 trials per condition, respectively), while emotion identification was tested using 6 × 15 and 12 × 20 configurations (120 trials per condition).

Human psychophysics and DNN evaluation used identical stimuli and conditions, ensuring direct comparability across agents.

### Computational observer models

We evaluated 14 DNN architectures spanning convolutional and transformer-based families, implemented using the timm library [86]. These included ConvNeXt models [75, 76], ResNets [74], VGG networks [73], Vision Transformers (ViT) [77], and a DINOv3 Vision Transformer [78], encompassing both convolutional and self-attention–based visual representations. All architectures were initialized from publicly available large-scale natural-image pretraining.

Models were trained under two transfer learning regimes. In the linear probe setting, pretrained backbones were frozen and only a task-specific linear classification head was optimized. In the end-to-end fine-tuning setting, all model parameters were updated jointly using differential learning rates for the pretrained backbone and the newly initialized classification head.

Training was performed using the AdamW optimizer [87] with a batch size of 32 for 30 epochs. For linear probe models, the classification head was trained with a learning rate of 10^−3^. For end-to-end fine-tuning, the classification head and backbone were trained with learning rates of 10^−4^ and 10^−5^, respectively, with weight decay of 0.01 applied in all cases. Architecture-specific input preprocessing was used throughout, implemented in PyTorch [88]. Each model was trained across 10 random seeds, with hyperparameters held constant across runs to ensure fair comparison between architectures and training regimes.

### Internal Noise Injection Procedures

We implemented two internal noise injection procedures in the end-to-end fine-tuned neural network classifiers to bring down performance closer to human levels. Both procedures use Monte Carlo estimation to approximate model accuracy under Gaussian noise added to model logits prior to the arg max operation [89, 59, 90].

#### Additive Noise Injection

In the additive noise procedure, we optimized a single noise parameter *σ* per task-architecture coupling to minimize the mean squared error (*MSE*) between human and model proportion correct (*PC*) across experimental conditions (i.e., implant device configurations). For each condition, we estimated model *PC* via Monte Carlo simulation. We trained *R* independent model replications (i.e., random seeds) per architecture. For each replication *r* and Monte Carlo run *n*, we drew a noise vector 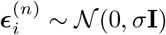 for each trial and computed proportion correct across all trials in condition *c*:

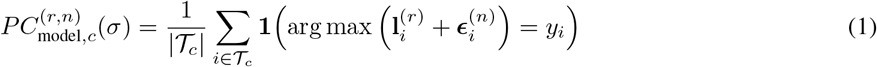

where 𝒯_*c*_ is the set of trials in condition *c* and 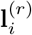 are the logits for trial *i* from replication *r*. The final model PC for condition *c* was obtained by averaging across *N* = 10,000 Monte Carlo runs and *R* replications:

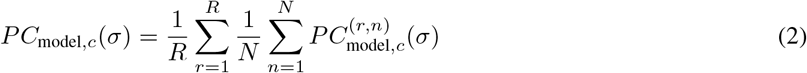

The optimal *σ* was determined by minimizing:

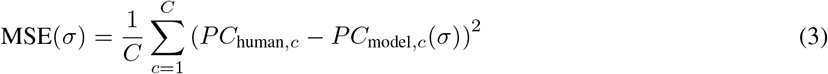

where *C* is the number of conditions and *PC*_human,*c*_ is the mean human accuracy for condition *c*.

#### Proportional Noise Injection

The proportional noise injection procedure used the same Monte Carlo estimation framework—including averaging across replications—but incorporated condition-specific baseline noise parameters *σ*_*c*_ derived from the variability of the trained model’s output logits, scaled by a global factor *a*.

To estimate *σ*_*c*_, we measured the dispersion of model logits across targets within each condition. For each task / architecture / device configuration, we grouped model outputs by target stimulus and computed the standard deviation of logits across trials for each output class, then averaged across classes and targets:

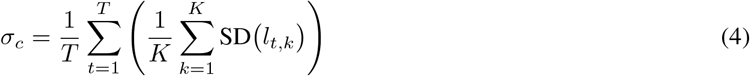

where *l*_*t,k*_ denotes the logit for class *k* and target *t, K* is the number of output classes, and *T* is the number of targets in the condition.

For each condition *c*, we drew 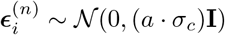, computing 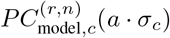 and averaging across Monte Carlo runs and replications as in Equations (1) and (2). The optimal scaling factor *a* was determined by minimizing:

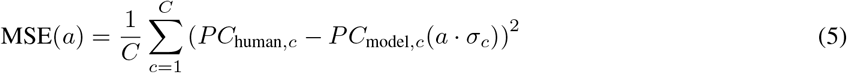

### Human-model alignment metrics

Model–human correspondence was quantified through multiple complementary performance metrics. We first computed proportion correct (*PC*), the raw classification accuracy representing the fraction of correctly classified trials. To account for task-specific chance performance (the shape task was unique in that it had 3 possible classes), we also calculated normalized above-chance proportion correct (*PC*^∗^) as

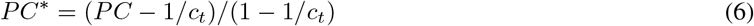

where *c*_*t*_ denotes the number of classes for task *t*. This normalization mapped chance performance to zero and perfect performance to one.

To quantify task-level consistency in performance trends across experimental conditions, we computed Pearson correlations between human and model accuracy across all shared implant configurations and phosphene models (*N* = 28). Standard errors were estimated via bootstrap resampling with *N* = 10,000 iterations, where subjects were sampled with replacement within each task, mean accuracy was recomputed by configuration, and the correlation coefficient was recalculated to yield an empirical distribution of correlation values under subject-level variability.

To assess whether models trained with linear probing achieved significantly higher human–model *PC* correlation than end-to-end and noise-injection baselines, we tested for differences in Pearson correlation coefficients using the bootstrap distributions described above. For each architecture and method pair, the *N* = 10,000 bootstrap iterations yielded an empirical distribution of correlation values under subject-level variability. Statistical significance was evaluated by computing the proportion of bootstrap samples in which the linear probe correlation fell below that of each comparison method (end-to-end fine-tuning, additive noise injection, and proportional noise injection), yielding a one-tailed bootstrap *p*-value. To control for multiple comparisons across architectures, *p*-values were adjusted using the Benjamini–Hochberg false discovery rate (FDR) procedure. Architecture-pooled (overall) *p*-values were computed by averaging the bootstrapped correlations across architectures prior to comparison, providing a summary statistic across the full model set for each bootstrap sample.

Beyond accuracy and correlation measures, we assessed the overlap in error patterns between humans and models using the Jaccard similarity index. For any two agents (human or model), the Jaccard index quantifies the proportion of shared errors relative to the total number of errors made by either agent, computed as

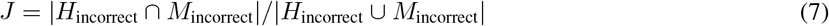

where *H*_incorrect_ and *M*_incorrect_ denote the sets of misclassified stimuli for agents *H* and *M*. Values approaching one indicate that both agents systematically fail on the same subset of stimuli, while values near zero suggest distinct error patterns.

As a reference baseline, we computed pairwise human–human Jaccard indices across all subject pairs and calculated a weighted mean based on the number of common trials per pair. Model–human Jaccard indices were computed analogously by pairing each model run with each human subject and averaging across all subject-run pairs weighted by trial count. Standard errors for both human–human and model–human Jaccard indices were estimated via subject-level bootstrap resampling with *N* = 10,000 iterations. To assess statistical significance, we performed permutation tests at the subject-run level. For each subject-run pair, we computed a null distribution of Jaccard indices by randomly shuffling model error labels *N* = 10,000 times while holding human errors fixed. The *p*-value was defined as the proportion of permuted Jaccard values meeting or exceeding the observed value, aggregated across all subject-run pairs using weighted averaging.

## Code and data availability

Code/modeling results available at https://github.com/jskaza/deep-learning-bionic-eyes.

## Acknowledgments

This research was supported by the UC Noyce Initiative, which fosters collaborative research in digital innovation across the University of California campuses. For more information, visit https://ucnoyce.org/. We thank Clara Yen and Adam Rychtecky for their invaluable assistance in coordinating and administering the psychophysics sessions.

## Supplementary Information

**Figure S1:**
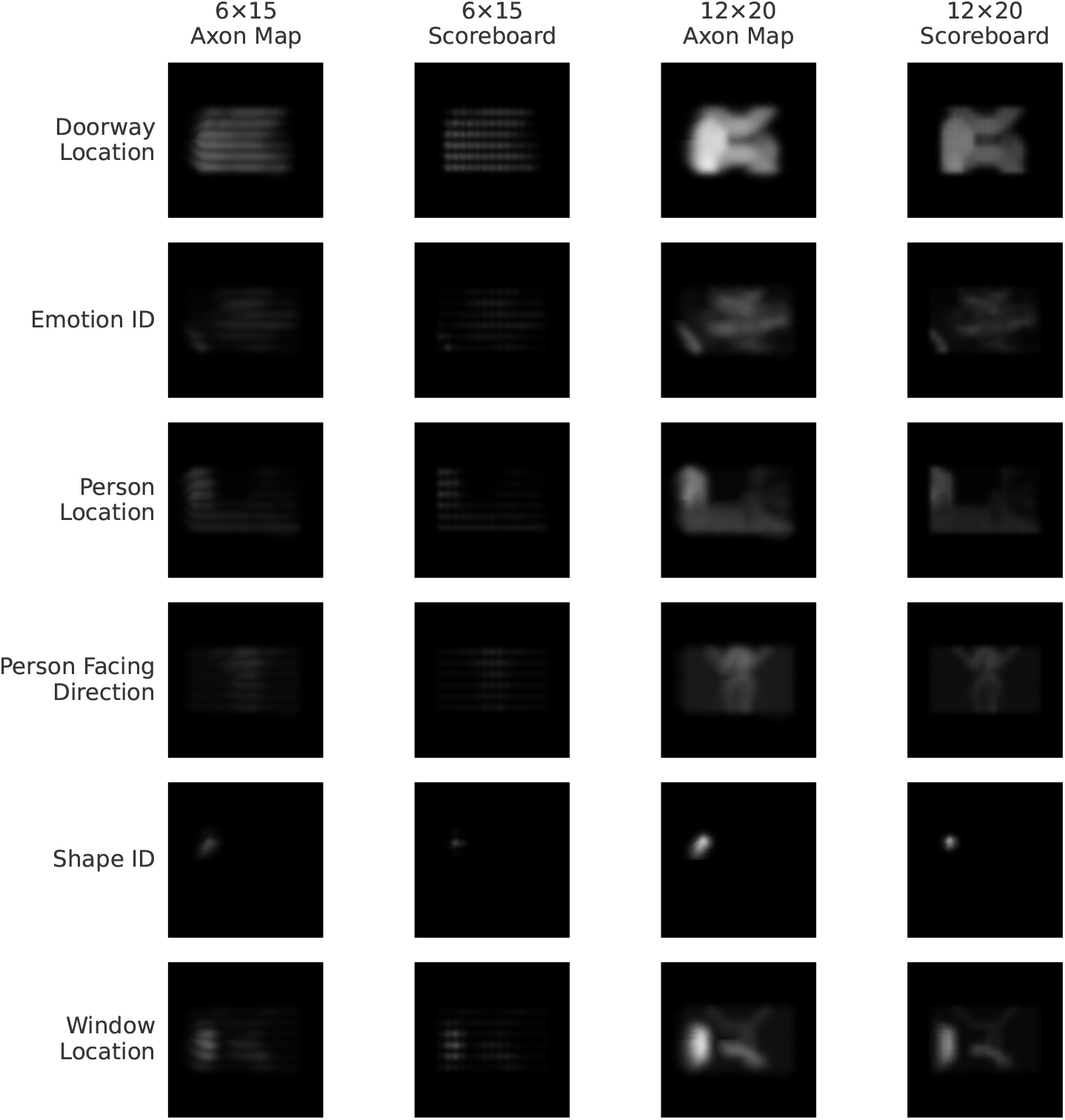
Example stimuli. Examples of phosphene stimuli for each task, both phosphene models, and the 6 × 15 and 12 × 20 electrode patterns.

**Table S1:** Model–human alignment for each architecture and training method, measured by Pearson correlation (*r*) and Jaccard similarity index (*J*). Pearson *r* reflects how well a model’s accuracy across 28 task–configuration combinations tracks human accuracy on the same conditions. Jaccard *J* reflects the trial-level overlap between errors made by the model and errors made by humans. Each row corresponds to one architecture trained with either linear probing (only the classification head is trained) or end-to-end fine-tuning (all weights are updated). Standard errors and p-values are based on bootstrap resampling of subjects. Models are sorted by Pearson *r* in descending order.

| Architecture | Method | Pearson $r$ | SE ($r$) | $p$ ($r$) | Jaccard Index ($J$) | SE ($J$) | $p$ ($J$) |
| --- | --- | --- | --- | --- | --- | --- | --- |
| VGG19 | Linear Probe | 0.941 | 0.018 | <0.001 | 0.169 | 0.013 | <0.001 |
| VGG11 | Linear Probe | 0.941 | 0.019 | <0.001 | 0.175 | 0.012 | <0.001 |
| VGG16 | Linear Probe | 0.940 | 0.019 | <0.001 | 0.169 | 0.013 | <0.001 |
| ResNet18 | Linear Probe | 0.938 | 0.018 | <0.001 | 0.212 | 0.011 | <0.001 |
| VGG13 | Linear Probe | 0.929 | 0.019 | <0.001 | 0.165 | 0.013 | <0.001 |
| ConvNeXt Small | Linear Probe | 0.916 | 0.016 | <0.001 | 0.149 | 0.007 | <0.001 |
| ResNet50 | Linear Probe | 0.911 | 0.023 | <0.001 | 0.199 | 0.011 | <0.001 |
| ViT Base | Linear Probe | 0.903 | 0.017 | <0.001 | 0.122 | 0.010 | <0.001 |
| ViT Large | Linear Probe | 0.901 | 0.016 | <0.001 | 0.124 | 0.010 | <0.001 |
| DINOv3 ViT Base | Linear Probe | 0.896 | 0.016 | <0.001 | 0.118 | 0.010 | <0.001 |
| ConvNeXt Tiny | Linear Probe | 0.893 | 0.015 | <0.001 | 0.162 | 0.008 | <0.001 |
| VGG13 | End-to-End | 0.883 | 0.021 | <0.001 | 0.123 | 0.010 | <0.001 |
| ConvNeXt Large | Linear Probe | 0.881 | 0.017 | <0.001 | 0.131 | 0.004 | <0.001 |
| VGG11 | End-to-End | 0.879 | 0.023 | <0.001 | 0.132 | 0.011 | <0.001 |
| ViT Small | Linear Probe | 0.877 | 0.018 | <0.001 | 0.124 | 0.010 | <0.001 |
| ConvNeXt Base | Linear Probe | 0.874 | 0.018 | <0.001 | 0.164 | 0.006 | <0.001 |
| DINOv3 ViT Base | End-to-End | 0.855 | 0.016 | <0.001 | 0.097 | 0.008 | <0.001 |
| VGG16 | End-to-End | 0.854 | 0.029 | <0.001 | 0.155 | 0.013 | <0.001 |
| VGG19 | End-to-End | 0.853 | 0.030 | <0.001 | 0.161 | 0.014 | <0.001 |
| ViT Small | End-to-End | 0.762 | 0.018 | <0.001 | 0.053 | 0.006 | <0.001 |
| ConvNeXt Base | End-to-End | 0.754 | 0.018 | <0.001 | 0.046 | 0.006 | <0.001 |
| ConvNeXt Large | End-to-End | 0.752 | 0.018 | <0.001 | 0.049 | 0.007 | <0.001 |
| ViT Base | End-to-End | 0.750 | 0.019 | <0.001 | 0.050 | 0.007 | <0.001 |
| ViT Large | End-to-End | 0.748 | 0.019 | <0.001 | 0.053 | 0.007 | <0.001 |
| ConvNeXt Small | End-to-End | 0.747 | 0.018 | <0.001 | 0.045 | 0.006 | <0.001 |
| ConvNeXt Tiny | End-to-End | 0.737 | 0.018 | <0.001 | 0.045 | 0.007 | <0.001 |
| ResNet18 | End-to-End | 0.719 | 0.038 | <0.001 | 0.195 | 0.017 | <0.001 |
| ResNet50 | End-to-End | 0.675 | 0.040 | <0.001 | 0.200 | 0.018 | <0.001 |

**Table S2:** Pearson correlation (*r*) between model and human accuracy across task conditions, for each architecture under two training methods and two internal noise paradigms. Linear Probe trains only a classification head on frozen features; End-to-End fine-tunes all model weights. Additive Noise and Proportional Noise replace learned representations with noise-driven responses, serving as fitted baselines. The Overall row reports the mean *r* across all architectures.

| Architecture | Linear Probe | End-to-End | Additive Noise | Proportional Noise |
| --- | --- | --- | --- | --- |
| ConvNeXt Base | 0.874 | 0.754 | 0.858 | 0.939 |
| ConvNeXt Large | 0.881 | 0.752 | 0.851 | 0.931 |
| ConvNeXt Small | 0.916 | 0.747 | 0.863 | 0.898 |
| ConvNeXt Tiny | 0.893 | 0.737 | 0.850 | 0.911 |
| DINOv3 ViT Base | 0.896 | 0.855 | 0.881 | 0.750 |
| ResNet18 | 0.938 | 0.719 | 0.865 | 0.841 |
| ResNet50 | 0.911 | 0.675 | 0.774 | 0.755 |
| VGG11 | 0.941 | 0.879 | 0.912 | 0.897 |
| VGG13 | 0.929 | 0.883 | 0.910 | 0.893 |
| VGG16 | 0.940 | 0.854 | 0.901 | 0.888 |
| VGG19 | 0.941 | 0.853 | 0.911 | 0.900 |
| ViT Base | 0.903 | 0.750 | 0.874 | 0.887 |
| ViT Large | 0.901 | 0.748 | 0.875 | 0.888 |
| ViT Small | 0.877 | 0.762 | 0.856 | 0.916 |
| **Overall** | **0.910** | **0.784** | **0.870** | **0.878** |

**Table S3:** Bootstrap p-values testing whether Linear Probe model–human correlation is significantly lower than that of alternative methods. Each comparison asks: does the Linear Probe correlate less well with human performance than End-to-End fine-tuning, End-to-End with Additive Noise, or End-to-End with Proportional Noise? P-values are estimated as the fraction of 10,000 subject-level bootstrap resamples in which the Linear Probe correlation fell below that of the comparison method, and are corrected for multiple comparisons.

| Architecture | Linear Probe < End-to-End (E2E) | Linear Probe < E2E + Add. | Linear Probe < E2E + Prop. |
| --- | --- | --- | --- |
| ConvNeXt Base | <0.001 | 0.085 | 1.000 |
| ConvNeXt Large | <0.001 | 0.006 | 1.000 |
| ConvNeXt Small | <0.001 | <0.001 | 0.050 |
| ConvNeXt Tiny | <0.001 | <0.001 | 1.000 |
| DINOv3 ViT Base | <0.001 | 0.042 | <0.001 |
| ResNet18 | <0.001 | <0.001 | <0.001 |
| ResNet50 | <0.001 | <0.001 | <0.001 |
| VGG11 | <0.001 | <0.001 | <0.001 |
| VGG13 | <0.001 | <0.001 | <0.001 |
| VGG16 | <0.001 | <0.001 | <0.001 |
| VGG19 | <0.001 | <0.001 | <0.001 |
| ViT Base | <0.001 | <0.001 | 0.017 |
| ViT Large | <0.001 | <0.001 | 0.046 |
| ViT Small | <0.001 | 0.003 | 1.000 |
| **Overall** | **<0.001** | **<0.001** | **<0.001** |

